# SECNVs: A Simulator of Copy Number Variants and Whole-Exome Sequences from Reference Genomes

**DOI:** 10.1101/824128

**Authors:** Yue Xing, Alan R. Dabney, Xiao Li, Guosong Wang, Clare A. Gill, Claudio Casola

## Abstract

Copy number variants are insertions and deletions of 1 kb or larger in a genome that play an important role in phenotypic changes and human disease. Many software applications have been developed to detect copy number variants using either whole-genome sequencing or whole-exome sequencing data. However, there is poor agreement in the results from these applications. Simulated datasets containing copy number variants allow comprehensive comparisons of the operating characteristics of existing and novel copy number variant detection methods. Several software applications have been developed to simulate copy number variants and other structural variants in whole-genome sequencing data. However, none of the applications reliably simulate copy number variants in whole-exome sequencing data. We have developed and tested SECNVs (Simulator of Exome Copy Number Variants), a fast, robust and customizable software application for simulating copy number variants and whole-exome sequences from a reference genome. SECNVs is easy to install, implements a wide range of commands to customize simulations, can output multiple samples at once, and incorporates a pipeline to output rearranged genomes, short reads and BAM files in a single command. Variants generated by SECNVs are detected with high sensitivity and precision by tools commonly used to detect copy number variants. SECNVs is publicly available at https://github.com/YJulyXing/SECNVs.

## 1 Introduction

Copy number variants (CNVs) represent DNA insertions and deletions ranging from a few dozen base pairs to several million bases that have been associated with phenotypic changes and human disease (Feuk et al., 2006). Initially discovered by array-based methods (Pinkel et al., 1998), CNVs have been increasingly detected using next-generation sequencing (NGS) data (Shen et al., 2019). A substantial proportion of CNVs encompass protein-coding genes (Zmienko et al., 2014). Many software applications have been developed to detect CNVs using either whole-genome sequencing (WGS) (Bartenhagen and Dugas, 2013; Pattnaik et al., 2014; Qin et al., 2015; Faust, 2017; Xia et al., 2017) or whole-exome sequencing (WES) (Sathirapongsasuti et al., 2011; Fromer et al., 2012; Klambauer et al., 2012; Koboldt et al., 2012a; Koboldt et al., 2012b; Krumm et al., 2012; Plagnol et al., 2012; Magi et al., 2013) data.

Whole-exome sequencing is based on the capture and sequencing of transcribed regions (exons) of protein coding sequences, which combined represent approximately 1% of the human genome. Thus, WES offers a significant benefit in terms of the sequencing costs compared to WGS. Additionally, WES data are an increasingly important source to identify genetic variants in non-model organisms (Lu et al., 2016; Kaur and Gaikwad, 2017). In species with very large genomes and limited opportunities for WGS experiments, WES data are expected to represent a critical source of information to detect CNVs (Hirsch et al., 2014).

Detection strategies for CNVs from next-generation sequencing data consist of four different approaches based on read depth, physical distance between read pairs (or paired-end mapping), detection of split reads, and comparison of de novo and reference genome assemblies (Alkan et al., 2011; Pirooznia et al., 2015). Because each of these approaches have limitations, programs that combine multiple strategies to detect CNVs based on WGS datasets have also been developed (see (Pirooznia et al., 2015)). In WES data, the approaches based on the distance between read pairs and detection of split reads have limited efficacy because the boundaries of the CNV region must fall completely within a target region for a CNV to be detected (Fromer et al., 2012; Alkodsi et al., 2014). However, the target regions only span a sparse 1% of the whole genome, therefore most of the breakpoints of CNVs are not located in the captured target regions (Tan et al., 2014; Yao et al., 2017). Thus, read depth represents the only effective strategy to detect CNVs from WES datasets and has been implemented in several programs (reviewed in (Tan et al., 2014)). Because these programs are built using different implementations and statistical models, they tend to produce datasets of CNVs with relatively little overlap (Magi et al., 2013; Kadalayil et al., 2014; Tan et al., 2014; Nam et al., 2016; Yao et al., 2017; Zare et al., 2017; Pounraja et al., 2019). Whole-exome sequencing data tend to have higher levels of noise and specific biases compared to WGS data (Zare et al., 2017), making detection of CNVs from WES data less accurate overall. In addition, there are limitations of the read depth method that make CNV detection in WES data less accurate (Tan et al., 2014). These limitations include poor resolution, systematic group effects, GC bias and difficulty in prediction of breakpoints in WES datasets (Tan et al., 2014). Therefore, benchmark analyses are necessary to evaluate the performance of CNV detection programs that utilize exome-sequencing datasets. Both simulated exome data and data from either arrays or WGS have enabled the assessment of CNV detection tools for WES datasets (Magi et al., 2013; Kadalayil et al., 2014; Tan et al., 2014; Nam et al., 2016; Yao et al., 2017; Zare et al., 2017). Simulations allow a more comprehensive assessment of the accuracy and power of these tools.

Most software applications developed to simulate CNVs fall short of generating the required outputs for WES datasets and are difficult to implement or cannot be applied to certain datasets (Table 1). Here, we introduce SECNVs (Simulator of Exome Copy Number Variants), a fast, robust and customizable software application for simulating CNV datasets using WES data. It relies upon a completely new approach to simulate test genomes and target regions to overcome some of the limitations of other WGS CNV simulation tools, and is the first ready-to-use WES CNV simulator. The simulator can be easily installed and used on Linux and MAC OS systems to facilitate comparison of the performance of different CNV detection methods and to test the most appropriate parameter settings for CNV identification.

**Table 1.** Comparison of current simulators to SECNVs.

| Simulator Type | Name | Language | Steps | Extra Files? * | Output Format | Short Reads Simulated? |
| --- | --- | --- | --- | --- | --- | --- |
| WGS | RSVSim<br>(Bartenhagen and Dugas, 2013) | R | Multiple | No | Test genome (fasta) and CNV information | No |
|  | SCNVSim<br>(Qin et al., 2015) | JAVA | 2 | Samtools index of reference genome; Chromosome length file; Repeat mask file | Test genome (fasta) and CNV information | No |
|  | Pysim-sv (Xia et al., 2017) | Python | Multiple | No | Fastq and SAM | Some |
|  | SVsim (Faust, 2017) | Python | 1 | Samtools index of reference genome | Test genome (fasta) and bedpe | No |
|  | SInC<br>(Pattnaik et al., 2014) | C | 2 | No | Test genome (fasta) and short reads | Some |
| WES | VarSimLab<sup>1</sup><br>(CNV-Sim<sup>2</sup>) | Python | 1 | No | Short reads or SAM, CNV information | Some |
|  | SECNVs | Python | 1 | No | Test genome (fasta), short reads and/or BAM files, and CNV information | Yes |

\* Extra files mean any files required in addition to the reference genome and target regions files.
<sup>1</sup> https://varsimlab.readthedocs.io/en/latest/index.html
<sup>2</sup> https://nabavilab.github.io/CNV-Sim/

## 2 Methods

### 2.1 Characteristics needed for a simulator of copy number variants

To generate WES reads, specific regions of a reference genome, called “target regions,” are captured and sequenced (Goh and Choi, 2012). To reproduce a realistic distribution of structural variants, a CNV simulator for WES data should generate variants that overlap partly or entirely with one or more target regions (Figure 1). The WES CNV detection tools require a list of target regions (exons) (Sathirapongsasuti et al., 2011; Fromer et al., 2012; Klambauer et al., 2012; Koboldt et al., 2012a; Koboldt et al., 2012b; Krumm et al., 2012; Plagnol et al., 2012; Magi et al., 2013), which can be obtained from public databases, and so this list could be used as the input to simulate short reads for those regions (Koboldt et al., 2009; Sathirapongsasuti et al., 2011; Koboldt et al., 2012a; Plagnol et al., 2012; Tan et al., 2014). Short reads would be simulated from a control genome (same as the reference genome) and test genomes (with simulated CNVs, SNPs, and indels) and aligned back to the control genome. In the read alignment file for the test genome, simulated CNVs (insertions and deletions) would ideally appear as increased read coverage or reduced read coverage, respectively (Figure 1). Options to generate short reads rearranged according to customized length, type (insertion or deletion) and copy number of CNVs within the genomic coordinates of target regions (whole exons and, potentially, regions upstream and downstream of exons) should be available in such a program (Figure 1). To mimic real data, it would be desirable to introduce SNPs and indels during this step as well.

**Figure 1.**
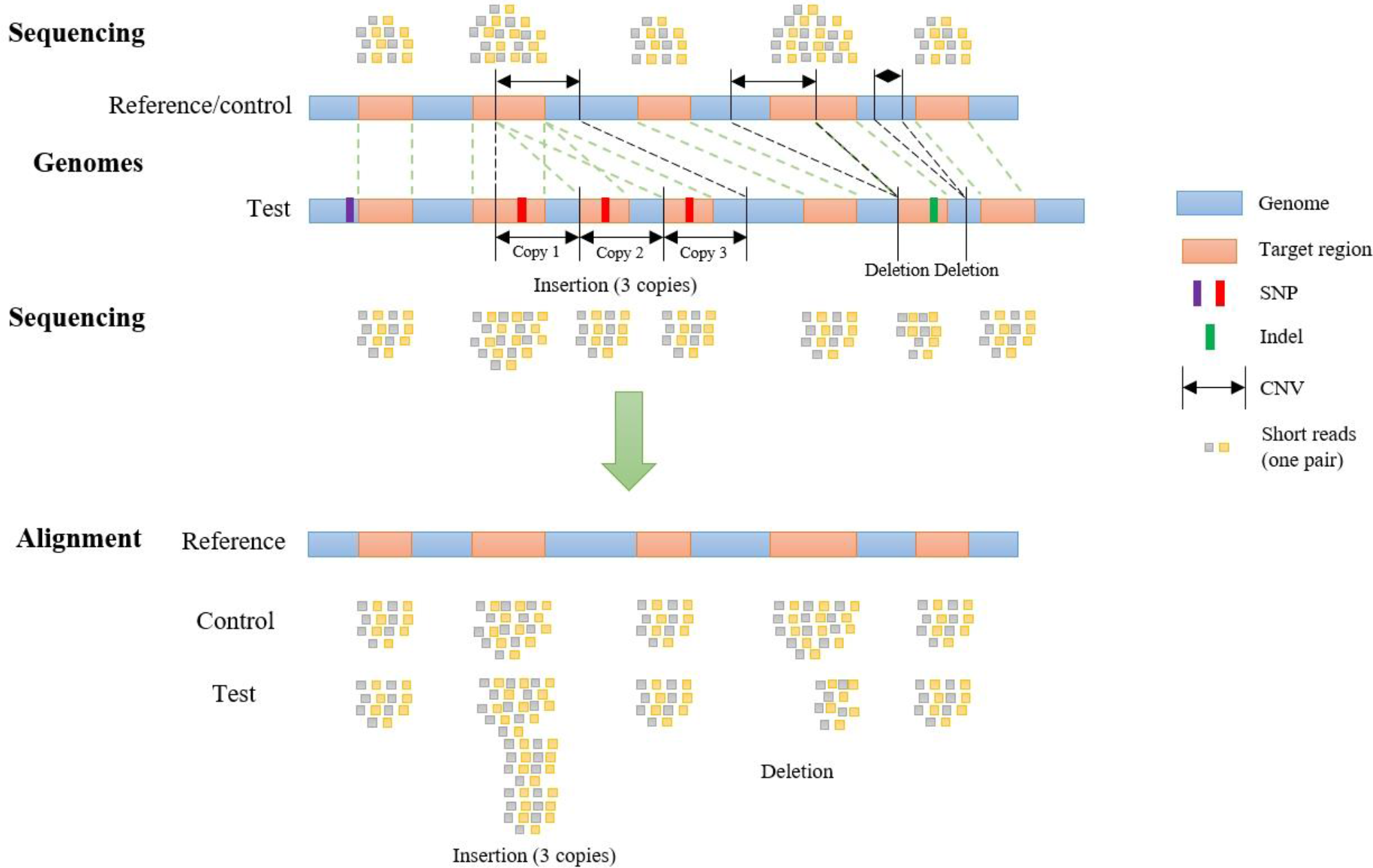
CNV detection by alignment of WES reads to a reference genome. WES data are obtained by sequencing target regions in genomes of interest. If the test genome contains insertions and/or deletions overlapping target regions, these regions will be rearranged (duplicated, deleted or shifted in their genomic coordinates) compared to control and reference genome. Reads from the test and control genomes are aligned to the original target regions in the reference genome. CNVs are detected according to the alignment.

These characteristics were incorporated into the program SECNVs, which we designed to solve the issue of how to reliably simulate CNVs for WES datasets. The Python-based SECNVs pipeline copies the FASTA reference genome (control) and a list of start and end coordinates for exons to a working directory. From the command line, the user can choose to expand or connect regions to specify targets for sequencing and define the type, total number, copy number, and length of CNVs to simulate. SECNVs makes a list of randomly generated CNVs and using that information creates a file of rearranged target regions. Next, FASTA-formatted test genome sequence(s) and FASTQ-formatted short sequence reads for the target regions from the control and single or pooled test genome(s) are simulated, and BAM file(s) and index(es) for them are all generated in a single command. These files can then be used as the input to compare various CNV detection tools.

### 2.2 Simulation of rearranged genomes and rearranged target regions

Before simulating WES, test genomes containing simulated CNVs that overlap with target regions are produced. First, the reference sequence is preprocessed based on how the user wants to handle gaps in the sequence. Next, a list of coordinates for CNVs are generated. Then SNPs and indels are simulated to create test genomes that mimic real data. Finally, CNVs are created in the FASTA-formatted test genome files.

#### 2.2.1 Preprocessing

First, SECNVs reads in a FASTA reference genome file and a file of target regions, and checks which option the user chose to handle ambiguous nucleotides (N) or assembly gaps (collectively referred to herein as ‘gaps’). An assembly gap is a stretch of 50 (default) or more “Ns” in the sequence. The user can choose to replace ambiguous nucleotides or gaps with random nucleotides, to avoid simulating CNVs in regions containing gaps, or to ignore the presence of gaps (default). If the user chose replacement, SECNVs finds gaps in the reference genome and fills them with random nucleotides. Instead, if the user chose to avoid them, after finding the gaps, SECNVs stores the genomic coordinates that demarcate each gap for the following steps.

#### 2.2.2 Creating a list of coordinates for copy number variant regions

Before actually simulating a FASTA test genome and WES reads that contain CNVs, a list of sites where the CNVs will be placed is generated by the software. Placement of CNVs in the sequence depends on many user defined parameters: proportion of each type (insertion, deletion), total number, range of copy number, range and distribution of lengths (random, Gaussian, Beta, user-supplied), spacing (random, Gaussian), and minimum spacing between CNVs. Unless the user specifies a number of CNVs per chromosome, the application considers the proportion of CNVs that would be expected on each chromosome based on the length of the chromosome. The software randomly allocates whether each CNV is an insertion or deletion and the number of copies will be simulated within the user- defined range for copy number, and the length is also assigned randomly within the user-defined range and length distribution. Once the length of the CNVs have been determined, for each CNV, the software randomly chooses the start point of that CNV based on CNV spacing and calculates the coordinate for the end point. At this stage, the software stores the coordinates for the beginning and end of the CNV region. Next, if the user specified that CNVs should not overlap with any gaps, SECNVs checks the coordinates of the CNV region against the coordinates for gaps. If an overlap is found, the CNV is discarded; otherwise it is kept for the next step. SECNVs then compares the start and end coordinate of the CNV region to the list of target regions. If there is partial (default minimum overlap is 50 bp) or complete overlap with targets, the region is retained, otherwise it is discarded, and the loop starts again. Before writing the coordinates for the CNV regions to the file, SECNVs checks for overlap with previously generated regions or a user-defined buffer region. Only non-overlapping CNV regions are recorded in the final list from SECNVs. The loop is repeated until the total number of CNVs is reached, unless the chromosome is too small and/or the number of target regions is too limited to simulate enough CNVs. In this situation, SECNVs outputs a warning message and the number of CNVs simulated on that chromosome is printed instead of the user specified number, and the program continues for other chromosomes. Users can also choose to simulate CNVs outside of target regions. The process is very similar to simulating CNVs overlapping with target regions. The only difference is that if a CNV doesn’t overlap with any target region, it will be kept; otherwise it will be discarded. SECNVs can also work with a list of predefined CNV regions. In this case SECNVs will read in the CNV list and use it as the final output of this step.

The Gaussian distribution of CNV spacing is generated by random selection from a symmetrically truncated Gaussian distribution mapped to the length of the chromosome, with distribution parameters (mean, SD) supplied by the user. Likewise, Gaussian distribution of CNV length is generated by random selection from a symmetrically truncated Gaussian distribution mapped to the range of user specified CNV length given distribution parameters (mean, SD) supplied by the user. The Beta distribution of CNV length, which is more realistic for CNV length distribution (Bartenhagen and Dugas, 2013), is generated by random selection from a Beta distribution mapped to the range of user-specified CNV lengths, again given distribution parameters (alpha, beta) from the user. Default values for alpha and beta are those used in Bartenhagen and Dugas 2013. Otherwise, the user must estimate the parameters for the Beta distribution using a collection of sample CNV lengths from their own data, which can easily be done using R (Team, 2016). Detailed instructions for this are included in the manual.

The final product of this step is a list of coordinates for CNV regions that are used in the following step to produce test genomes that are rearranged from the reference genome and adjusted coordinates for target regions.

#### 2.2.3 Simulation of test genomes and adjusted target regions

The target regions are duplicated, deleted or shifted as a result of the simulated CNVs, as shown in Figure 1. Short reads are generated based on the rearranged genome and target regions, and aligned to the original reference/control genome in this “simulation of short reads” step.

Before introducing CNVs into the FASTA genome sequence, SNPs and indels are simulated as requested by the user. First, SNPs are randomly generated in the target regions plus a user-supplied buffer region upstream and downstream of the target regions (default is 0), based on the SNP rate specified by the user. In this step, SECNVs randomly extracts n positions from these regions to simulate SNPs, where n equals the total length of the regions multiplied by the SNP rate. Then, nucleotides for these positions are randomly changed to another nucleotide in the test genome using the weights assigned by (Park, 2009) to represent the known mutation rate for SNP in human. Users can modify the mutation rates for other organisms. Detailed instructions are in the manual.

Next, indels are randomly generated in the target regions based on the indel rate (default is 0) specified by the user. In this step, m start points of indels are randomly generated in the target regions, where m equals the total length of the target regions multiplied by the indel rate. The length of each indel is then assigned by randomly choosing a number between 1 and the maximum indel length specified by the user. Type of indel (insertion or deletion) is randomly assigned to each indel as well. Next, SECNVs sorts the indels by their start points, and generates them one by one. If an indel is an insertion, SECNVs will make a random string of nucleotides of the previously assigned length and insert it at the assigned start point of that indel in the test genome sequence. Then, SECNVs recalculates the genomic coordinates of the target regions. If the start and/or end of the target regions are greater than the start point of the indel, their coordinates are increased by the length of that indel. The start point of the remaining indels is iteratively changed as well: coordinates of subsequent indels are increased by the length of that indel.

If an indel is a deletion, SECNVs will first check if the length of that indel is smaller than the target region it is in. If not, the length of that indel is reduced to ensure that at least one base pair of the target region remains. Then the sequence between the coordinates defining the indel is deleted from the test genome. Next, if the start and/or end of the target regions are greater than the start point of the indel, their coordinates are adjusted by subtracting the length of that indel. The start point of the rest of the indels are adjusted in the same way.

Finally, after the SNP and indels are created, the simulated CNVs are generated in the FASTA test genome files. In general, users would simulate CNVs that overlap with targets. The list of CNVs is sorted by coordinate and processed one by one so that the coordinates for subsequent CNVs are adjusted, similar to the process used for indels.

For each CNV, the genomic start and end coordinates and length are extracted. Then SECNVs loops iteratively through the genomic coordinates for all the target regions on a chromosome. If a target region is completely inside the CNV, it is categorized as “inside the CNV.” If a target region partially overlaps with a CNV, it will be split into at least two parts: the parts outside of the CNV and the part overlapping with the CNV, and then categorized (upstream, inside, downstream), as shown in Figure 2. Sometimes users will choose to simulate CNV completely outside of target regions, even though those CNV will be undetectable in the WES. If a target region is completely before the CNV, it is categorized as “upstream of the CNV” and if a target region is completely after the CNV, it is categorized as “downstream of the CNV.”

**Figure 2.**
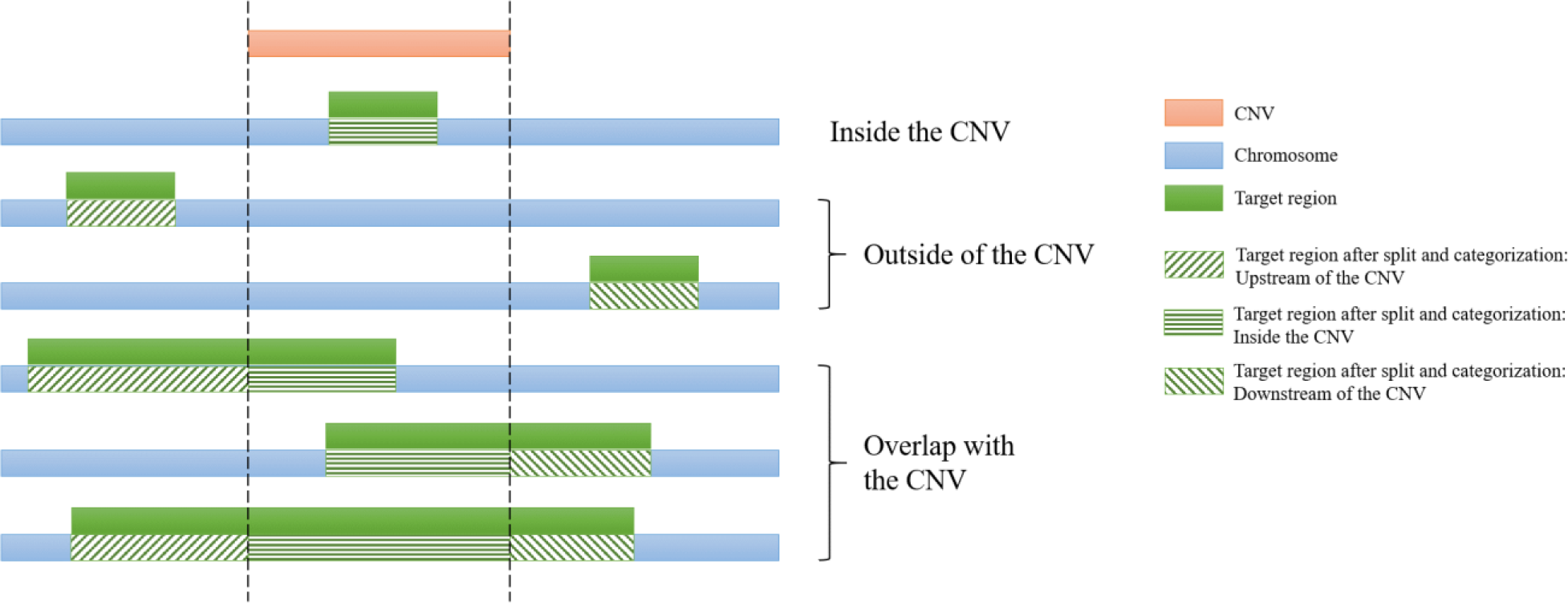
Categorization of target regions in the test genome. For each of the simulated CNVs, all target regions on a chromosome are assigned as “upstream of the CNV,” “inside the CNV,” and/or “downstream of the CNV”. If the region partially overlaps it is also split. Afterwards genomic coordinates for the targets are recalculated and split regions are reconnected.

The next step is to adjust the coordinates to take into account the placement of the CNV relative to the target regions. As shown in Supplementary Figure 1, the coordinates of target regions categorized as “upstream of the CNV” remain unchanged. Coordinates of target regions categorized as “inside the CNV” must be adjusted. For insertions, the new start and end positions of these target regions will be: new position = length ∗ (number of copies − 1) + old position, where the number of copies loops from 1 to the total copy number of that insertion, thus creating a tandemly duplicated CNV insertion event in the test sequence. For deletions, when the CNV and target region overlap, the coordinates for that part of the target are deleted from the file. Coordinates of target regions categorized as “downstream of the CNV” will be altered as follows. For insertions, new position = length ∗ (total copy number of the insertion − 1) + old position. For deletions, new position = old position − length of the CNV.

Finally, the CNV sequence will be copied to or deleted from the FASTA test genome accordingly. All the genomic coordinates of CNVs subsequent to this CNV in the list are adjusted in the same manner as the target regions categorized as “downstream of the CNV.” Finally, if a target region was previously split, it will be reconnected.

The software loops iteratively through all CNVs to create the rearranged test genome sequence and target regions for short read simulation. If the user chose to simulate multiple test genomes, the steps after preprocessing will be repeated to simulate each test genome.

The output files generated by this step include: 1. Test genome(s) (FASTA) with non-overlapping CNVs; 2. Target regions for test genome(s) (.bed); 3. Control genome (FASTA, optional); 4. Target regions for control genome (.bed, always generated in case the target regions are modified by user in sequencing steps); 5. List(s) of CNVs overlapping with target regions (.bed); 6. List(s) of CNVs outside of target regions (.bed, optional).

This is the core step of SECNVs. Users can choose to continue the pipeline within SECNVs to simulate short reads or use another short read simulator and the files produced from this step as input.

### 2.3 Simulation of short reads

Users have the option to generate short read files with SECNVs by simulating single- or paired-end sequences from the test and control genomes. During this step, if the spacing between target regions is less than the spacing selected by user (default 0), the target regions are connected to form a single region (called a combined target region) to simulate the sequences. Users can also choose to expand the target regions by including additional nucleotides (default 0) upstream and downstream of the target regions (called an extended target region) for sequencing. The number of reads, type of reads (paired-end or single-end), fragment size, standard deviation of fragment size, read length, quality score offset, and error model can also be specified. A default error model: Illumina HighSeq 2500 for WES paired end sequencing is provided. This default error model was generated using a modified GemSIM script (McElroy et al., 2012) which fixed a bug to make the error profile generation function work. The dataset used for generating this error model was a human WES dataset from the Sequence Read Archive at the National Center for Biotechnology Information: run number ERR3385637. Users can also generate their own error model from real data using this modified GemSIM script, to keep the error profiles up to date as sequencing technology changes over time. Detailed instructions on how to use it to make new error profiles are included in the manual.

Instead, SECNVs reads in the headers of the input file as keys of a dictionary and reads the sequences line by line and combines them as values of that dictionary for the corresponding keys. Short read sequences are generated within the combined and extended target regions, which match just the target regions when default settings are used. Reads passing GC filtering are synthesized using a modification to the Wessim1 (Kim et al., 2013) algorithm (ideal target approach). Wessim1 only simulates reads at the start and the end of each target region (Supplementary File 1). Custom codes were written to modify Wessim1’s scripts to correct this shortcoming of the program. Now fragments across the entire target regions based on fragment size and standard deviation of fragment size are produced and saved as FASTQ sequence, better mimicking real-world WES sequencing data. Output files from this step are the short reads for test genome(s) (FASTQ) and the short reads for the control genome (FASTQ, optional).

### 2.4 Creating BAM files and indexes from the simulated short read files

BAM files and indexes can be generated from the short read files for the test and control genomes through a standard pipeline that implements the widely-used tools BWA (Li, 2013), samtools (Li et al., 2009; Li, 2011), Picard^3^ and GATK (McKenna et al., 2010):

1. The Burrows-Wheeler Aligner of BWA is used to align the FASTQ reads to create a SAM file.
2. Samtools is used to convert the file format to a BAM file, sort the BAM file, and remove potential PCR duplicates.
3. Picard is used to add read groups to the samples.
4. GATK is used to locally realign reads, to minimize the number of mismatching bases across all the reads.

The output files in this step include: 1. Indexes for the control genome (.dict, .fai, .sa, etc., if BAM files are to be generated and no indexes exist in the output directory); 2. BAM file(s) and index(es) for the test genome(s) (.bam and .bai); 3. BAM file(s) and index(es) for control genome (.bam and .bai, optional).

### 2.5 Validation of method

To confirm that the code for the algorithm implemented in SECNVs is correctly simulating the test genome and target regions, a small pseudo-genome was used as input to illustrate the process in Supplementary Figure 1.

### 2.6 Example command lines

1. Simulate 10 CNVs overlapping with target regions, and 1 CNV outside of target regions randomly on each chromosome using default lengths, copy numbers, minimum distance between each of the 2 CNVs and proportion of insertions. For each CNV overlapping with target regions, the overlapping length is no less than 90 bps. CNV break points follow a Gaussian(1, 2) distribution, and CNV lengths follow a Beta(2, 5) distribution. CNVs are not generated in gaps. A total of 5 test and control samples are built. Short reads (fastq) files are generated using default settings, paired-end sequencing. SECNVs/SECNVs.py -G <input_fasta> -T <target_region> -o <output_dir> \ -e_chr 10 -o_chr 1 -ol 90 -ms gauss -as 1 -bs 2 -ml beta -al 2 -bl 5 -eN gap -n 5 -sc -pr -ssr
2. Simulate CNVs overlapping with target regions from a provided CNV list. Twenty CNVs are to be simulated outside of target regions randomly on the whole genome with default settings. CNVs are not to be generated on any stretches of “N”s. A pair of test and control genome are built. SECNVs/SECNVs.py -G <input_fasta> -T <target_region> -o <output_dir> \ -e_cnv <list_of_CNV_overlapping_with_target_regions> -o_tol 20 -eN all -sc
3. Simulate 20 CNVs overlapping with target regions on the whole genome, and have at least 100 bps between any 2 CNVs. CNVs are not generated outside of target regions. Gaps (50 or more consecutive “N”s) are replaced by random nucleotides. SNP rate is 0.001 and indel rate is 0.00001, and the maximum indel length is 100 bps. Paired-end sequencing reads with quality offset 35 are then produced. For a pair of test and control genomes BAM files are generated. SECNVs/SECNVs.py -G <input_fasta> -T <target_region> -o <output_dir> \ -e_tol 20 -f 100 -rN gap -sc -pr -q 35 -ssr -sb \ -s_r 0.001 -i_r 0.00001 -i_mlen 100 \ -picard <absolute_path_to_picard> -GATK <absolute_path_to_GATK>
4. Simulate CNVs overlapping with target regions and outside of target regions from provided files of CNV lengths. Combined single regions are formed from two or more regions originally separated by less than 100 bps. CNVs are not generated on gaps (60 or more consecutive “N”s). A total of 10 test and control samples are built. The paired-end sequencing must include sequences 50 bp upstream and downstream of the target regions. The final output consists of short reads (fastq) files with 100,000 reads. SECNVs/SECNVs.py -G <input_fasta> -T <target_region> -o <output_dir> \ -ml user -e_cl <length_file_1> -o_cl <length_file_2> \ -clr 100 -eN gap -n_gap 60 -n 10 -sc -pr -tf 50 -nr 100000 -ssr

### 2.7 Simulation of mouse and human WES datasets

To evaluate the performance of SECNVs, we used mouse (mm10) and human (hg38) chromosome 1 (downloaded from UCSC genome browser: https://genome.ucsc.edu/) as control genomes. Target regions were exons, which were also downloaded from the UCSC genome browser. We simulated 20 test genomes for each species that included 100 randomly distributed CNVs that overlapped at least 100 bp of target regions and ranged from 1,000 to 100,000 bp in length. Another 10 CNVs outside of target regions were also generated for each species. For each test genome, all sequences with “Ns” (gaps) were excluded, the SNP rate was set at 10^−3^, and the indel rate was set at 10^−5^(Mills et al., 2006). The minimum distance between any two CNVs was 1000 bp. For the synthesis of short reads, target regions less than 100 bp apart were connected and 50 bp upstream and downstream of the connected target regions were also sequenced. Paired-end sequencing was set to a base quality offset of 33. A total of 1 million reads were generated for each sample, with a fragment size of 200 bp and a read length of 100 bp. The rearranged fasta genome files with target regions, fastq short read files, BAM files and indexes for the 20 samples and control were simulated in one command for each species as follows:

Mouse: python SECNVs/SECNVs.py -G mouse/mouse.1.fa -T mouse/mouse.1.bed -e_chr 100 - o_chr 10 -o test_mous -rn mouse -ssr -sb -f 1000 -ol 100 -tf 50 -clr 100 -sc -eN all -pr -n 20 -q 33 - s_r 0.001 -i_r 0.00001 -nr 1000000 -picard <absolute_path_to_picard> -GATK <absolute_path_to_GATK>
Human: python SECNVs /SECNVs.py -G hg38/hg38.1.fa -T hg38/hg38.1.bed -e_chr 100 -o_chr 10 - o test_human_nn -rn human -ssr -sb -f 1000 -ol 100 -tf 50 -clr 100 -sc -eN all -pr -n 20 -q 33 -s_r 0.001 -i_r 0.00001 -nr 1000000 -picard <absolute_path_to_picard> -GATK <absolute_path_to_GATK>

### 2.8 Validation of read generation and alignment

To confirm SECNVs is reliably simulating short reads and BAM files, read alignments in the target regions for human and mouse chromosome 1 against the respective reference genome were visualized with IGV (Thorvaldsdóttir et al., 2013).

### 2.9 Evaluation of performance of SECNVs

To demonstrate the utility of simulated datasets generated by SECNVs in performance testing of CNV detection software, we chose three commonly used WES CNV detection tools: ExomeDepth (Plagnol et al., 2012), CODEX2 (Jiang et al., 2018), and CANOES (Backenroth et al., 2014). Performance was evaluated for sensitivity, precision and false discovery rate. Sensitivity is the number of true CNVs that are correctly detected divided by the total number of true CNVs. Precision is the number of CNVs correctly detected by tools, divided by the total number of CNVs detected by tools. False discovery rate (FDR), which equals to 1 – precision, is the number of CNVs incorrectly detected by tools, divided by the total number of CNVs detected by tools. During this evaluation, we found that CNV transition probability in ExomeDepth and CNV occurrence in CANOES influenced the test results the most, so we evaluated different values for these parameters as well. All other parameters were either left as default or set to fit the characteristics of CNVs we expected to detect. For example, in ExomeDepth, length of expected CNVs was set to 50000, which was about the average CNV length expected. A CNV was considered detected if at least 80% of the detected CNV overlap with a simulated CNV. We also used the best application with optimized parameter settings to test if any CNVs outside of target regions were detected.

## 3 Results

In this study, we presented a fast, reliable and highly-customizable software application, SECNVs, which takes in a reference genome and target regions to simulate SNPs, indels and CNVs in one or multiple test genomes, as well as the control, and outputs fasta formatted genome files with target regions, short read files, BAM files and indexes in a single command.

### 3.2 Computational speed

SECNVs is a fast software application for CNV simulation. The detailed approximate computation time to generate CNVs for each sample on human and mouse chromosome 1 is shown in Table 2.

**Table 2.** Computation time and memory usage of SECNVs.

|  | Mouse chromosome 1 | Human chromosome 1 |
| --- | --- | --- |
| 1. Read in genome and target region files, exclude all “N” sequences | < 15 seconds | < 25 seconds |
| 2. Generate list of CNVs overlapping with target regions | < 4 min | < 8 min |
| 3. Generate list of CNVs outside of target regions | < 50 s | < 3 min |
| 4. Generate rearranged genome: make SNPs, indels and CNVs in the genome | < 50 s | < 80 s |
| 5. Generate short reads | < 7.5 min | < 7.5 min |
| 6. Create indexes for the control genome | < 3.5 min | < 4.5 min |
| 7. Generate BAM file and index | < 2 min | < 2 min |
| Max memory | 5630 MB | 6516 MB |
| Average memory | 1885.78 MB | 2521.26 MB |

### 3.3 Validation of method

Simulation of SNPs, indels and CNVs in the tiny pseudo-genome established that the method of random replacement of gaps, simulation of SNPs, indels and CNVs in the test genome is accurate; and the rearrangement of target regions for the test genome is accurate as well (Supplementary File 2; Supplementary Figure 1).

The simulated short reads and BAM files generated using the modified Wessim1 code align across the whole target regions (Figure 3). Differences in read coverage at combined and extended target regions in test and control BAM files are characteristic of CNVs spanning these target regions (Figure 3 a and c). For target regions without CNVs, there was no obvious difference in read coverage at combined and extended target regions in test and control BAM files (Figure 3 b and d). No reads were aligned outside of target regions, regardless of whether there were CNVs or not, except for a few alignment errors. The reliability of the BAM files ensured that CNVs overlapping with target regions could be readily detected.

**Figure 3.**
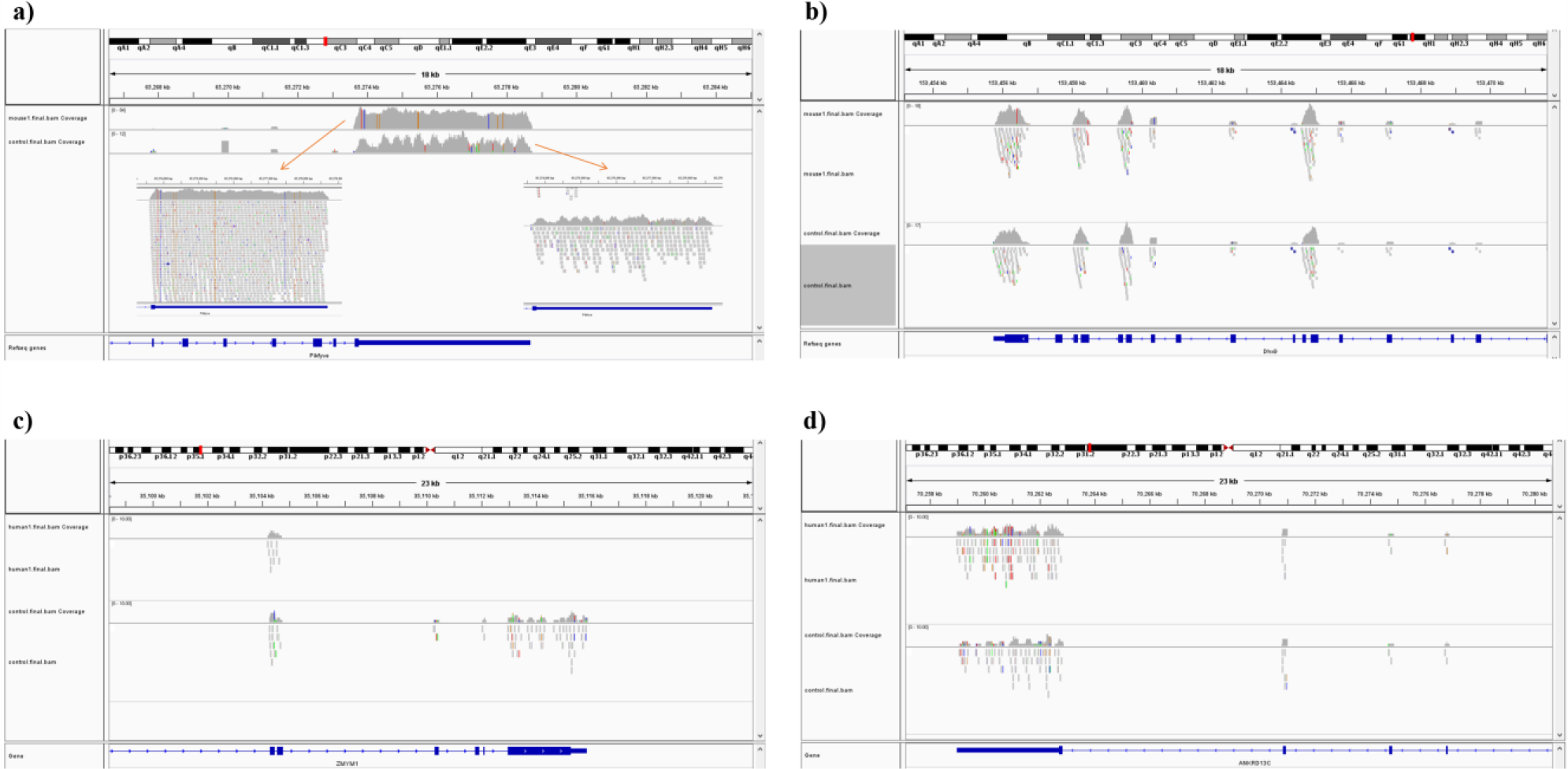
Exemplar simulated output BAM files visualized in IGV. a) A 10 copy insertion at mouse chr1:65272798-65339955, which partially overlap with exons of the Pikfyve gene. Because of the read depth in this region, the reads tracks are shown side-by-side; b) A region of Dhx9 gene of the mouse genome, showing no CNVs in this region; c) A deletion at human chr1: 35106158-35150376, which partially overlap with exons of the Zmym1 gene; d) A region of ANKRD13C gene of the human genome, showing no CNVs in this region. In each image, the top track is a region of the test genome and the middle track shows the same region of the control genome. The bottom track is the exons and introns of genes.

### 3.4 Sensitivity and precision of CNV detection from simulated WES datasets

Average sensitivity, precision, FDR and the number of CNVs detected by ExomeDepth, CANOES and CODEX2 using simulated reads from human and mouse genomes are summarized in Table 3 and Figure 4. Regardless of the species, as the transition probability from ExomeDepth increased (Figure 4 a and b) and until the number of CNV detected matched the number of CNV simulated, sensitivity increased, and precision was high. Beyond 100 CNV, the number of detected CNV rapidly inflated and precision rapidly declined. A similar profile was observed for occurrence of CNV in CANOES (Figure 4 c and d). The overall performance of ExomeDepth in terms of precision and sensitivity was better than CANOES or CODEX (Figure 5). False discovery rate was higher for the human data than the mouse data. CODEX was not able to detect all of the simulated CNV. Although the parameters of transition probability in ExomeDepth and occurrence of CNV in CANOES are similar in concept, the comparison in Figure 5 shows that as each of these parameters was changed, performance of the two software tools was very different. We also confirmed that CNVs simulated outside of exomes were never detected.

**Table 3.**
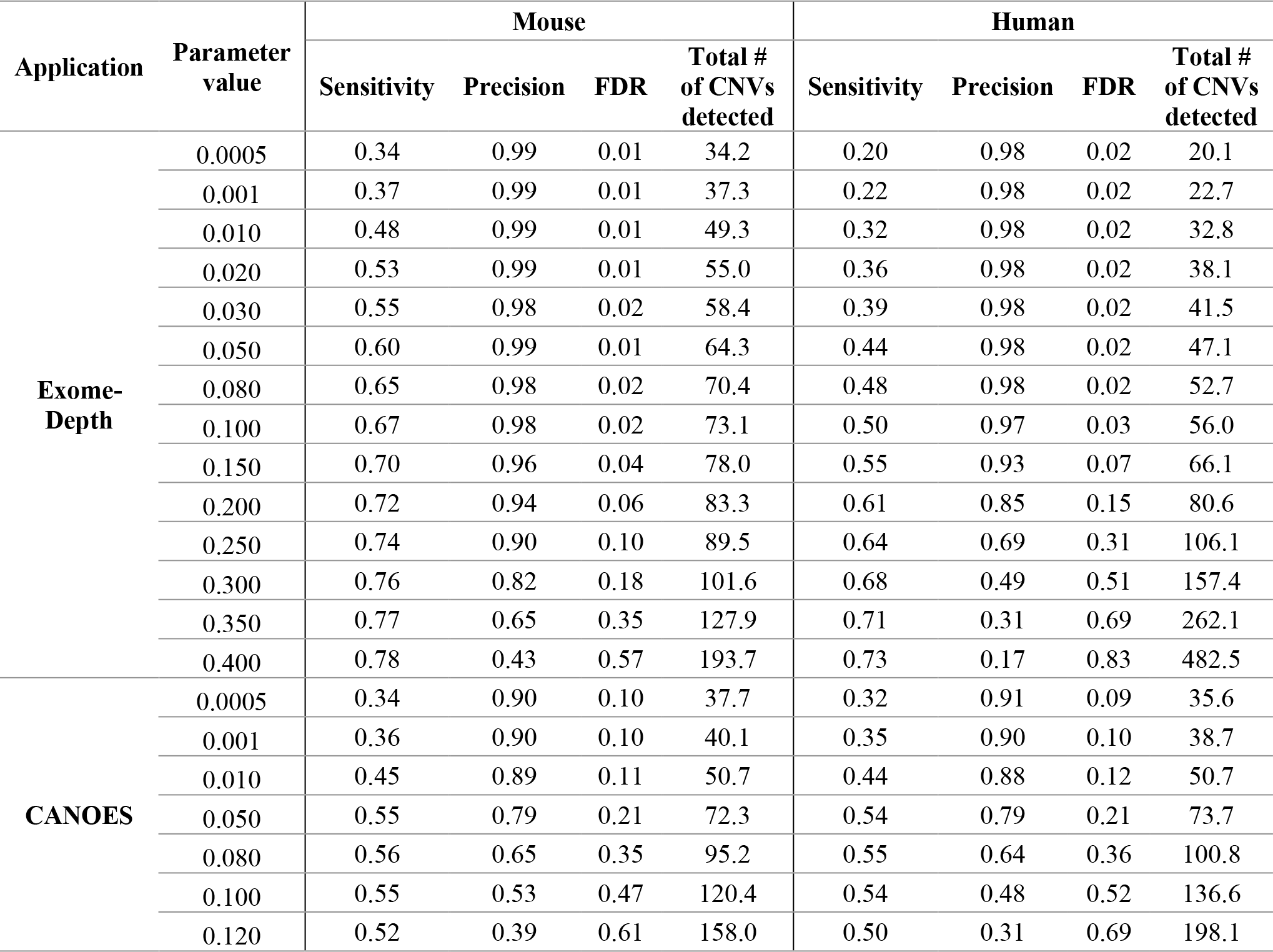
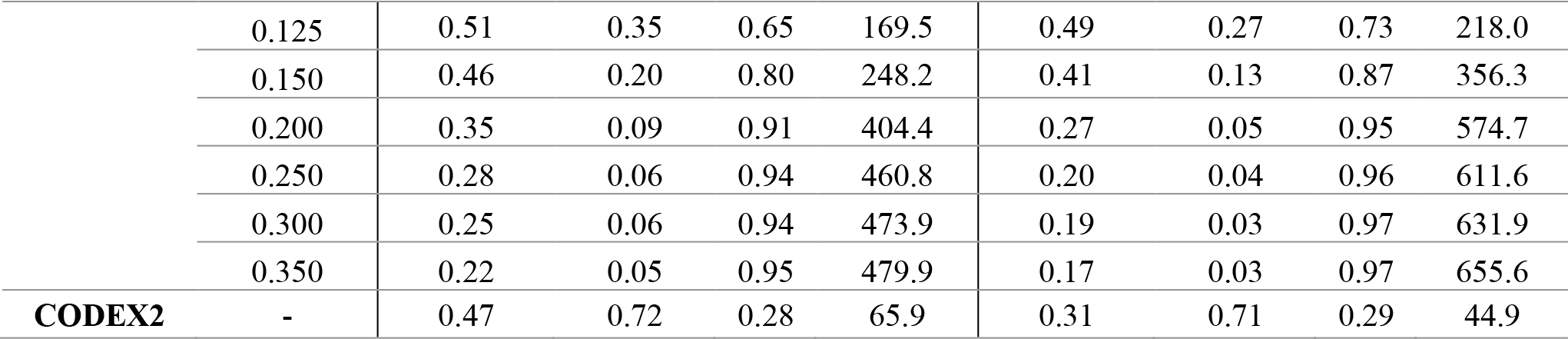
Average sensitivity, precision, FDR and number of CNV detected using simulated WES datasets. The parameter for ExomeDepth is transition probability and the parameter for CANOES is CNV occurrence.

| Application | Parameter value | Mouse |  |  |  | Human |  |  |  |
| --- | --- | --- | --- | --- | --- | --- | --- | --- | --- |
|  |  | Sensitivity | Precision | FDR | Total # of CNVs detected | Sensitivity | Precision | FDR | Total # of CNVs detected |
| Exome-Depth | 0.0005 | 0.34 | 0.99 | 0.01 | 34.2 | 0.20 | 0.98 | 0.02 | 20.1 |
|  | 0.001 | 0.37 | 0.99 | 0.01 | 37.3 | 0.22 | 0.98 | 0.02 | 22.7 |
|  | 0.010 | 0.48 | 0.99 | 0.01 | 49.3 | 0.32 | 0.98 | 0.02 | 32.8 |
|  | 0.020 | 0.53 | 0.99 | 0.01 | 55.0 | 0.36 | 0.98 | 0.02 | 38.1 |
|  | 0.030 | 0.55 | 0.98 | 0.02 | 58.4 | 0.39 | 0.98 | 0.02 | 41.5 |
|  | 0.050 | 0.60 | 0.99 | 0.01 | 64.3 | 0.44 | 0.98 | 0.02 | 47.1 |
|  | 0.080 | 0.65 | 0.98 | 0.02 | 70.4 | 0.48 | 0.98 | 0.02 | 52.7 |
|  | 0.100 | 0.67 | 0.98 | 0.02 | 73.1 | 0.50 | 0.97 | 0.03 | 56.0 |
|  | 0.150 | 0.70 | 0.96 | 0.04 | 78.0 | 0.55 | 0.93 | 0.07 | 66.1 |
|  | 0.200 | 0.72 | 0.94 | 0.06 | 83.3 | 0.61 | 0.85 | 0.15 | 80.6 |
|  | 0.250 | 0.74 | 0.90 | 0.10 | 89.5 | 0.64 | 0.69 | 0.31 | 106.1 |
|  | 0.300 | 0.76 | 0.82 | 0.18 | 101.6 | 0.68 | 0.49 | 0.51 | 157.4 |
|  | 0.350 | 0.77 | 0.65 | 0.35 | 127.9 | 0.71 | 0.31 | 0.69 | 262.1 |
|  | 0.400 | 0.78 | 0.43 | 0.57 | 193.7 | 0.73 | 0.17 | 0.83 | 482.5 |
| CANOES | 0.0005 | 0.34 | 0.90 | 0.10 | 37.7 | 0.32 | 0.91 | 0.09 | 35.6 |
|  | 0.001 | 0.36 | 0.90 | 0.10 | 40.1 | 0.35 | 0.90 | 0.10 | 38.7 |
|  | 0.010 | 0.45 | 0.89 | 0.11 | 50.7 | 0.44 | 0.88 | 0.12 | 50.7 |
|  | 0.050 | 0.55 | 0.79 | 0.21 | 72.3 | 0.54 | 0.79 | 0.21 | 73.7 |
|  | 0.080 | 0.56 | 0.65 | 0.35 | 95.2 | 0.55 | 0.64 | 0.36 | 100.8 |
|  | 0.100 | 0.55 | 0.53 | 0.47 | 120.4 | 0.54 | 0.48 | 0.52 | 136.6 |
|  | 0.120 | 0.52 | 0.39 | 0.61 | 158.0 | 0.50 | 0.31 | 0.69 | 198.1 |
|  | 0.125 | 0.51 | 0.35 | 0.65 | 169.5 | 0.49 | 0.27 | 0.73 | 218.0 |
|  | 0.150 | 0.46 | 0.20 | 0.80 | 248.2 | 0.41 | 0.13 | 0.87 | 356.3 |
|  | 0.200 | 0.35 | 0.09 | 0.91 | 404.4 | 0.27 | 0.05 | 0.95 | 574.7 |
|  | 0.250 | 0.28 | 0.06 | 0.94 | 460.8 | 0.20 | 0.04 | 0.96 | 611.6 |
|  | 0.300 | 0.25 | 0.06 | 0.94 | 473.9 | 0.19 | 0.03 | 0.97 | 631.9 |
|  | 0.350 | 0.22 | 0.05 | 0.95 | 479.9 | 0.17 | 0.03 | 0.97 | 655.6 |
| **CODEX2** | - | 0.47 | 0.72 | 0.28 | 65.9 | 0.31 | 0.71 | 0.29 | 44.9 |

**Figure 4.**
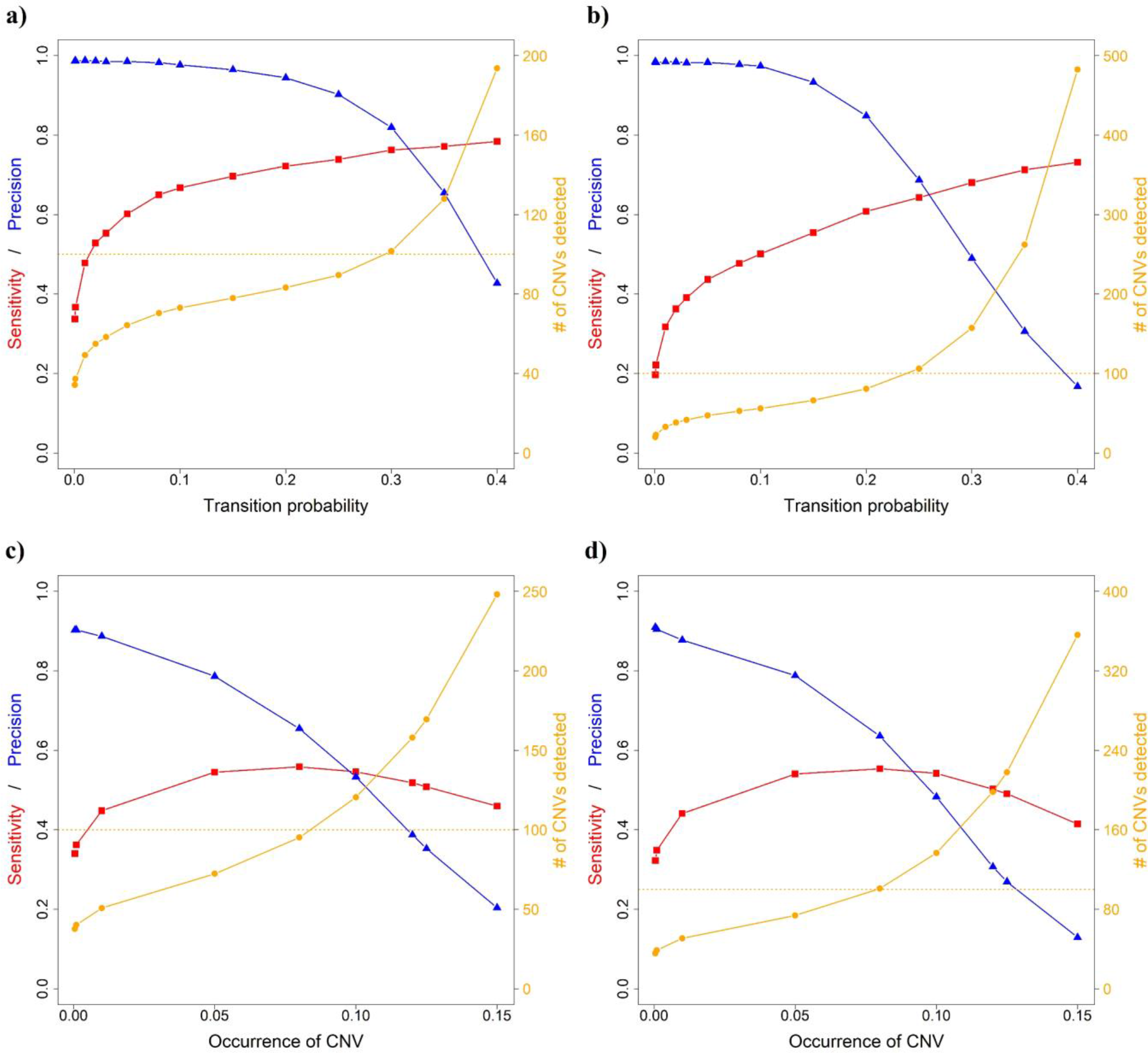
Sensitivity, precision and number of CNVs detected for ExomeDepth and CANOES. The sensitivity, precision and number of CNVs detected in a) simulated mouse data for ExomeDepth, b) simulated human data for ExomeDepth, c) simulated mouse data for CANOES and d) simulated human data for CANOES are displayed. Red lines show sensitivity, blue lines show precision and orange lines show the number of CNVs detected. Solid triangles, squares and circles represent the actual data points.

**Figure 5.**
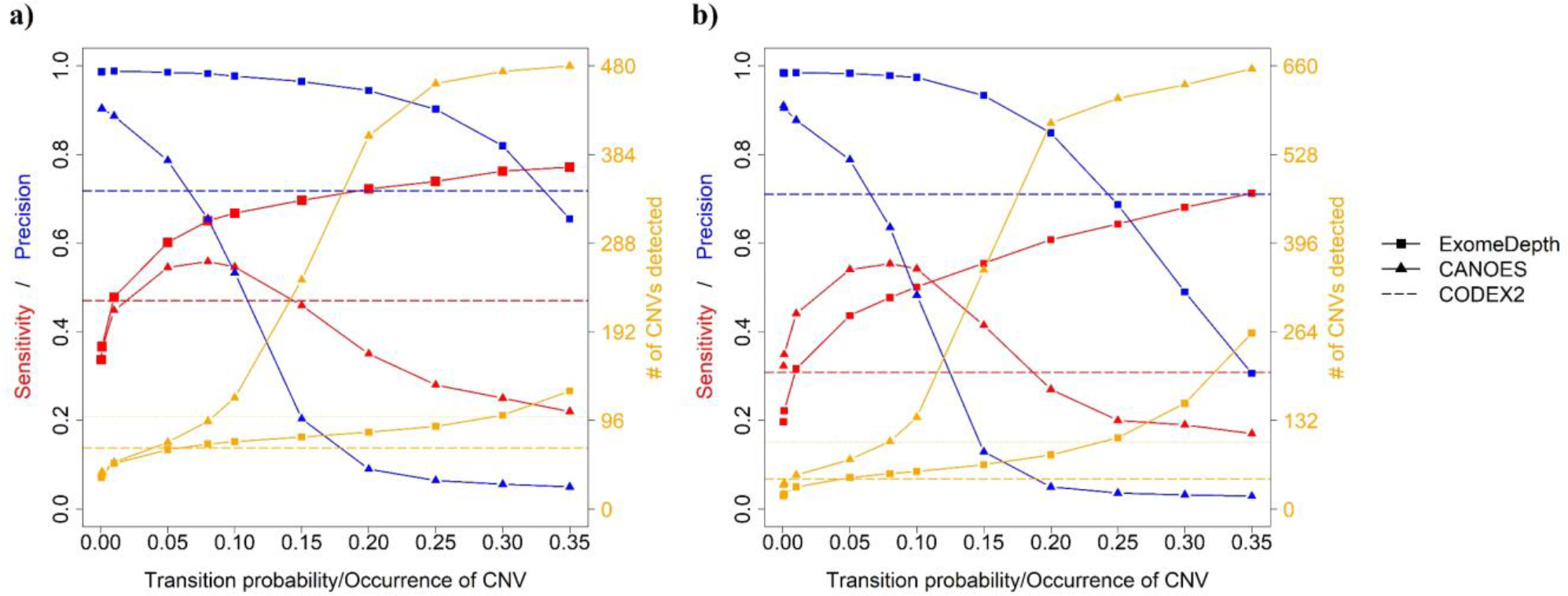
Comparison of sensitivity, precision and number of CNVs detected by the three software applications. a) simulated mouse data and b) simulated human data. Red lines show sensitivity, blue lines show precision and orange lines show the number of CNVs detected. Because CODEX2 does not have the parameter “transition probability” or “CNV occurrence,” a single value for sensitivity, precision and number of CNVs is displayed.

## 4 Discussion

Copy number variants represent an important source of genetic variation and have been associated with disease and other important phenotypic traits in humans, domesticated animals and crops [1; Zhang et al 2009; Alkan et al. 2011]. Whole-exome sequencing projects represent an increasingly common source of genomic data that can be harvested to detect CNVs. Often applied in the detection of mutations associated with cancer, Mendelian and complex diseases in humans, WES data have also been generated for multiple non-model organisms to identify genetic variants, including CNVs (Prunier et al., 2017; Low et al., 2019). However, previous studies have shown that detection of CNVs from WES data is inconsistent across the tools designed to detect these variants (Guo et al., 2013; Nam et al., 2016; Yao et al., 2017). These evaluations have largely relied on datasets of well-characterized CNVs obtained using array-based experiments or WGS data from human samples, a benchmarking approach that presents several limitations. First, known variants tend to occupy the higher end of the spectrum of lengths for CNVs. Second, known CNVs are often derived from cancer tissues and are expected to show different features than germline CNVs. Third, the characteristics of CNVs might differ significantly between humans and other organisms. Therefore, flexible CNV simulators would allow more rigorous testing of the efficacy of these tools.

Previously developed simulators fall short of producing realistic CNVs and present a variety of operational issues that make them challenging or impossible to use. Most of the applications for simulating CNVs and other structural variants from WGS data (Table 1) (Bartenhagen and Dugas, 2013; Pattnaik et al., 2014; Qin et al., 2015; Faust, 2017; Xia et al., 2017), require commands be entered in several steps, and require further processing to use their outputs. Among them, only RSVSim (Bartenhagen and Dugas, 2013) and SVsim (Faust, 2017) allow users to specifically generate CNVs in user-selected regions of the genome. However, they cannot calculate rearranged coordinates of the target regions in the test genome after simulating CNVs, which makes it impossible to use their outputs to generate accurate sets of short reads if the user only knows the original target regions but not the probe sequences in sequencing step. Additionally, RSVSim runs into an infinite loop when there are too many gaps in the genome.

To the best of our knowledge, VarSimLab^1^, previously released as CNV-Sim^2^, is the only other program specifically designed to simulate CNVs from WES data (Zare et al., 2017), but the website for the software indicates it is currently not usable. We found that CNV-Sim had a problem generating short reads. The main issue with these programs appears to be their reliance on the ideal target approach implemented in the application Wessim1 (Kim et al., 2013) to generate short reads, but this approach does not provide coverage across target regions (see Supplementary File 1). Furthermore, the majority of CNVs simulated by CNV-Sim overlap with each other when target regions are nearby on the reference genome. A third issue is that this program creates temporary “genomes” by deleting the segments between target regions. Additional copies for the test genome are generated in the case of insertions, and additional copies for the control genome are created in the case of deletions. Short read files are then generated using these test and control genomes (called “tumor” and “normal” in CNV-Sim). This makes it impossible to generate pooled samples with a common control, whereas many CNV detection applications for WES data require pooled samples as input. In addition, it cannot simulate the realistic scenario of CNVs with different degrees of overlap with target regions. Finally, CNV-Sim only accepts one chromosome at a time.

Here, we described SECNVs, a novel software application that fills this gap by simulating realistic CNVs from WES data. First, it uses a completely new method to accurately and reliably simulate test genome(s) and target regions with SNPs, indels and CNVs. Second, it incorporates a modified version of the Wessim1 algorithm to simulate short reads, which effectively mimics real-world WES sequencing, including GC filtration. Third, to keep the sequencing error profile up to date, SECNVs provides a recent error profile for short read simulation and includes detailed instructions on how to make user-specified error profiles from real data. Finally, the options for CNV simulation are highly customizable. In this paper, SECNVs was applied to human and mouse data and the results showed that CNVs simulated by the software application were successfully detected by various WES CNV detection software applications, demonstrating that output from SECNVs can be used to test these applications and their parameters.

Sensitivity and FDR were similar to previous reports using real data (Seiser and Innocenti, 2015; Zare et al., 2017). It is known that the sensitivity is low and FDR is high for CNV detection in WES datasets (Tan et al., 2014; Yao et al., 2017). This is because read depth approaches are the only reliable method for WES CNV detection, but they have many limitations (Tan et al., 2014). In addition, the high SNP and indel rate introduced, as well as GC filtration and sequencing errors affect the alignment and reduce sensitivity and increase FDR. However, compared to the real human CNV detection from Illumina genotyping microarrays by other Hidden Markov Model-Based CNV detection methods mentioned in (Seiser and Innocenti, 2015), and real human CNV detection from WES data using various software applications (Zare et al., 2017), the sensitivity, precision, and FDR all suggest that CNVs generated by SECNVs are reliable and can be easily detected.

After reaching the true CNV number, the number of detected CNVs tends to inflate. When this happens, sensitivity either increases very slowly or begins to drop, and precision decreases rapidly. Therefore, the user can choose the transition probability, CNV occurrence or similar parameter to detect CNVs approximating the number of real CNVs, and determine if the sensitivity and precision are acceptable. Alternatively, the user can sacrifice some sensitivity and detect fewer CNVs to get higher precision. Of course, users can test other parameters as well to find out the most suitable software application and parameter settings for their data.

Using datasets simulated with SECNVs, we were also able to characterize the performance of CNV detection software applications under a wide range of parameters. We showed that simulations are critical to assess the effect of key parameters on the sensitivity and accuracy of such applications. Thus, SECNVs can simulate highly customized WES datasets to mimic real-world data and enable users to identify the most appropriate software application and parameter settings for their real data.

We identified two main limitations in the current version of SECNVs. First, because SNPs and indels are simulated in the test genome and then CNVs are simulated, there is no variability among the duplicated sequences. Second, all CNVs are tandem duplications or deletions in SECNVs. However, these limitations do not affect CNV detection from WES data, because most WES CNV detection methods are read depth based, which cannot distinguish between tandem duplication and insertion elsewhere in the genome (Tan et al., 2014). The nature of WES datasets makes methods other than read depth ineffective for WES CNV detection (Tan et al., 2014).

One caution is that most diploid reference genomes report a consensus sequence for each pair of chromosomes. Therefore, by default the SECNVs simulator adds or deletes two copies at a time. If the user wanted to extend the simulator to an odd number of insertion or deletion events, the bam files for the reference and test genomes could be merged.

Currently, SECNVs only simulates indels in target regions to increase speed. For SNPs, buffer regions upstream and downstream of target regions are allowed for simulation, because SNPs downstream of CNVs may affect the detection of that CNV (Bartenhagen and Dugas, 2013). Buffer regions upstream and downstream of target regions are also allowed for short read simulation. In future version of SECNVs, SNPs and indels will be simulated for the whole genome.

The time used for each step implemented in SECNVs strongly depends on the parameter settings supplied by the user and increases in an approximately linear manner. For instance, run time is positively correlated with the number of CNVs, SNPs and indels that are generated. In addition, the run time tends to increase significantly when the length of all CNVs combined exceeds the length of the genome, but we expect that this model will rarely be implemented when simulating realistic CNVs even in small genomes. Given its flexibility, precision and variety of unique features, SECNVs represents a reliable application to study copy number variants using WES data for various species and under a variety of conditions.

## Author Contributions

YX designed, created and tested the software application, evaluated its performance, wrote the draft of the manuscript and contributed to its major revisions. ARD, CC and CAG provided suggestions on the design and update of the software application, contributed to major revisions of the manuscript, and read and approved the final manuscript. XL participated in design and test of the software application, and contributed to revisions of the manuscript. GW provided suggestions on the update of the software application and solved technical problems.

## Acknowledgments

The high-performance research computing system Ada at Texas A&M University was used to test the software application.

## Data Availability Statement

The genome sequence and target region (exon) files analyzed for this study can be found in the UCSC genome database at https://genome.ucsc.edu/cgi-bin/hgGateway (Consortium, 2001). The datasets simulated and analyzed during the current study are available from the corresponding author on request.

## Footnotes

1 https://varsimlab.readthedocs.io/en/latest/index.html

2 https://nabavilab.github.io/CNV-Sim/

3 https://broadinstitute.github.io/picard/

## References

Alkan, C., Coe, B.P., and Eichler, E.E. (2011). Genome structural variation discovery and genotyping. Nat Rev Genet 12(5), 363–376. doi: 10.1038/nrg2958.

Alkodsi, A., Louhimo, R., and Hautaniemi, S. (2014). Comparative analysis of methods for identifying somatic copy number alterations from deep sequencing data. Briefings in Bioinformatics 16(2), 242–254. doi: 10.1093/bib/bbu004.

Backenroth, D., Homsy, J., Murillo, L.R., Glessner, J., Lin, E., Brueckner, M., et al. (2014). CANOES: detecting rare copy number variants from whole exome sequencing data. Nucleic acids research 42(12), e97–e97. doi: 10.1093/nar/gku345.

Bartenhagen, C., and Dugas, M. (2013). RSVSim: an R/Bioconductor package for the simulation of structural variations. Bioinformatics 29(13), 1679–1681.

Consortium, I.H.G.S. (2001). Initial sequencing and analysis of the human genome. Nature 409(6822), 860.

Faust, G. (2017). “SVsim: a tool that generates synthetic Structural Variant calls as benchmarks to test/evaluate SV calling pipelines”.).

Feuk, L., Carson, A.R., and Scherer, S.W. (2006). Structural variation in the human genome. Nature Reviews Genetics 7(2), 85.

Fromer, M., Moran, J.L., Chambert, K., Banks, E., Bergen, S.E., Ruderfer, D.M., et al. (2012). Discovery and statistical genotyping of copy-number variation from whole-exome sequencing depth. American journal of human genetics 91(4), 597–607. doi: 10.1016/j.ajhg.2012.08.005.

Goh, G., and Choi, M. (2012). Application of whole exome sequencing to identify disease-causing variants in inherited human diseases. Genomics & informatics 10(4), 214–219.

Guo, Y., Sheng, Q., Samuels, D.C., Lehmann, B., Bauer, J.A., Pietenpol, J., et al. (2013). Comparative study of exome copy number variation estimation tools using array comparative genomic hybridization as control. BioMed research international 2013.

Hirsch, C.D., Evans, J., Buell, C.R., and Hirsch, C.N. (2014). Reduced representation approaches to interrogate genome diversity in large repetitive plant genomes. Briefings in Functional Genomics 13(4), 257–267. doi: 10.1093/bfgp/elt051.

Jiang, Y., Wang, R., Urrutia, E., Anastopoulos, I.N., Nathanson, K.L., and Zhang, N.R. (2018). CODEX2: full-spectrum copy number variation detection by high-throughput DNA sequencing. Genome Biology 19(1), 202. doi: 10.1186/s13059-018-1578-y.

Kadalayil, L., Rafiq, S., Rose-Zerilli, M.J.J., Pengelly, R.J., Parker, H., Oscier, D., et al. (2014). Exome sequence read depth methods for identifying copy number changes. Briefings in Bioinformatics 16(3), 380–392. doi: 10.1093/bib/bbu027.

Kaur, P., and Gaikwad, K. (2017). From Genomes to GENE-omes: Exome Sequencing Concept and Applications in Crop Improvement. Frontiers in plant science 8, 2164–2164. doi: 10.3389/fpls.2017.02164.

Kim, S., Jeong, K., and Bafna, V. (2013). Wessim: a whole-exome sequencing simulator based on in silico exome capture. Bioinformatics 29(8), 1076–1077.

Klambauer, G., Schwarzbauer, K., Mayr, A., Clevert, D.A., Mitterecker, A., Bodenhofer, U., et al. (2012). cn.MOPS: mixture of Poissons for discovering copy number variations in next-generation sequencing data with a low false discovery rate. Nucleic Acids Res 40(9), e69. doi: 10.1093/nar/gks003.

Koboldt, D.C., Chen, K., Wylie, T., Larson, D.E., McLellan, M.D., Mardis, E.R., et al. (2009). VarScan: variant detection in massively parallel sequencing of individual and pooled samples. Bioinformatics (Oxford, England) 25(17), 2283–2285. doi: 10.1093/bioinformatics/btp373.

Koboldt, D.C., Zhang, Q., Larson, D.E., Shen, D., McLellan, M.D., Lin, L., et al. (2012a). VarScan 2: somatic mutation and copy number alteration discovery in cancer by exome sequencing. Genome research 22(3), 568–576. doi: 10.1101/gr.129684.111.

Koboldt, D.C., Zhang, Q., Larson, D.E., Shen, D., McLellan, M.D., Lin, L., et al. (2012b). VarScan 2: somatic mutation and copy number alteration discovery in cancer by exome sequencing. Genome Res 22(3), 568–576. doi: 10.1101/gr.129684.111.

Krumm, N., Sudmant, P.H., Ko, A., O’Roak, B.J., Malig, M., Coe, B.P., et al. (2012). Copy number variation detection and genotyping from exome sequence data. Genome research 22(8), 1525–1532. doi: 10.1101/gr.138115.112.

Li, H. (2011). A statistical framework for SNP calling, mutation discovery, association mapping and population genetical parameter estimation from sequencing data. Bioinformatics 27(21), 2987–2993.

Li, H. (2013). Aligning sequence reads, clone sequences and assembly contigs with BWA-MEM. arXiv preprint arXiv:1303.3997.

Li, H., Handsaker, B., Wysoker, A., Fennell, T., Ruan, J., Homer, N., et al. (2009). The sequence alignment/map format and SAMtools. Bioinformatics 25(16), 2078–2079.

Low, T.Y., Mohtar, M.A., Ang, M.Y., and Jamal, R. (2019). Connecting Proteomics to Next-Generation Sequencing: Proteogenomics and Its Current Applications in Biology. Proteomics 19(10), e1800235. doi: 10.1002/pmic.201800235.

Lu, M., Krutovsky, K.V., Nelson, C.D., Koralewski, T.E., Byram, T.D., and Loopstra, C.A. (2016). Exome genotyping, linkage disequilibrium and population structure in loblolly pine (Pinus taeda L.). BMC Genomics 17(1), 730. doi: 10.1186/s12864-016-3081-8.

Magi, A., Tattini, L., Cifola, I., D’Aurizio, R., Benelli, M., Mangano, E., et al. (2013). EXCAVATOR: detecting copy number variants from whole-exome sequencing data. Genome Biology 14(10), R120. doi: 10.1186/gb-2013-14-10-r120.

McElroy, K.E., Luciani, F., and Thomas, T. (2012). GemSIM: general, error-model based simulator of next-generation sequencing data. BMC Genomics 13(1), 74. doi: 10.1186/1471-2164-13-74.

McKenna, A., Hanna, M., Banks, E., Sivachenko, A., Cibulskis, K., Kernytsky, A., et al. (2010). The Genome Analysis Toolkit: a MapReduce framework for analyzing next-generation DNA sequencing data. Genome research.

Mills, R.E., Luttig, C.T., Larkins, C.E., Beauchamp, A., Tsui, C., Pittard, W.S., et al. (2006). An initial map of insertion and deletion (INDEL) variation in the human genome. Genome research 16(9), 1182–1190. doi: 10.1101/gr.4565806.

Nam, J.-Y., Kim, N.K.D., Kim, S.C., Joung, J.-G., Xi, R., Lee, S., et al. (2016). Evaluation of somatic copy number estimation tools for whole-exome sequencing data. Briefings in bioinformatics 17(2), 185–192. doi: 10.1093/bib/bbv055.

Park, L. (2009). Relative mutation rates of each nucleotide for another estimated from allele frequency spectra at human gene loci. Genet Res (Camb) 91(4), 293–303. doi: 10.1017/s0016672309990164.

Pattnaik, S., Gupta, S., Rao, A.A., and Panda, B. (2014). SInC: an accurate and fast error-model based simulator for SNPs, Indels and CNVs coupled with a read generator for short-read sequence data. BMC bioinformatics 15(1), 40.

Pinkel, D., Segraves, R., Sudar, D., Clark, S., Poole, I., Kowbel, D., et al. (1998). High resolution analysis of DNA copy number variation using comparative genomic hybridization to microarrays. Nat Genet 20(2), 207–211. doi: 10.1038/2524.

Pirooznia, M., Goes, F.S., and Zandi, P.P. (2015). Whole-genome CNV analysis: advances in computational approaches. Front Genet 6, 138. doi: 10.3389/fgene.2015.00138.

Plagnol, V., Curtis, J., Epstein, M., Mok, K.Y., Stebbings, E., Grigoriadou, S., et al. (2012). A robust model for read count data in exome sequencing experiments and implications for copy number variant calling. Bioinformatics (Oxford, England) 28(21), 2747–2754. doi: 10.1093/bioinformatics/bts526.

Pounraja, V.K., Jayakar, G., Jensen, M., Kelkar, N., and Girirajan, S. (2019). A machine-learning approach for accurate detection of copy number variants from exome sequencing. Genome Res 29(7), 1134–1143. doi: 10.1101/gr.245928.118.

Prunier, J., Caron, S., and MacKay, J. (2017). CNVs into the wild: screening the genomes of conifer trees (Picea spp.) reveals fewer gene copy number variations in hybrids and links to adaptation. BMC Genomics 18(1), 97. doi: 10.1186/s12864-016-3458-8.

Qin, M., Liu, B., Conroy, J.M., Morrison, C.D., Hu, Q., Cheng, Y., et al. (2015). SCNVSim: somatic copy number variation and structure variation simulator. BMC bioinformatics 16(1), 66.

Sathirapongsasuti, J.F., Lee, H., Horst, B.A.J., Brunner, G., Cochran, A.J., Binder, S., et al. (2011). Exome sequencing-based copy-number variation and loss of heterozygosity detection: ExomeCNV. Bioinformatics 27(19), 2648–2654. doi: 10.1093/bioinformatics/btr462.

Seiser, E.L., and Innocenti, F. (2015). Hidden Markov Model-Based CNV Detection Algorithms for Illumina Genotyping Microarrays. Cancer informatics 13(Suppl 7), 77–83. doi: 10.4137/CIN.S16345.

Shen, W., Szankasi, P., Durtschi, J., Kelley, T.W., and Xu, X. (2019). Genome-Wide Copy Number Variation Detection Using NGS: Data Analysis and Interpretation. Methods Mol Biol 1908, 113–124. doi: 10.1007/978-1-4939-9004-7_8.

Tan, R., Wang, Y., Kleinstein, S.E., Liu, Y., Zhu, X., Guo, H., et al. (2014). An Evaluation of Copy Number Variation Detection Tools from Whole-Exome Sequencing Data. Human Mutation 35(7), 899–907. doi: 10.1002/humu.22537.

Team, R.C. (2016). “R: A Language and Environment for Statistical Computing”. (Vienna, Austria: R Foundation for Statistical Computing).

Thorvaldsdóttir, H., Robinson, J.T., and Mesirov, J.P. (2013). Integrative Genomics Viewer (IGV): high – performance genomics data visualization and exploration. Briefings in bioinformatics 14(2), 178–192.

Xia, Y., Liu, Y., Deng, M., and Xi, R. (2017). Pysim-sv: a package for simulating structural variation data with GC-biases. BMC bioinformatics 18(3), 53.

Yao, R., Zhang, C., Yu, T., Li, N., Hu, X., Wang, X., et al. (2017). Evaluation of three read-depth based CNV detection tools using whole-exome sequencing data. Molecular cytogenetics 10, 30–30. doi: 10.1186/s13039-017-0333-5.

Zare, F., Dow, M., Monteleone, N., Hosny, A., and Nabavi, S. (2017). An evaluation of copy number variation detection tools for cancer using whole exome sequencing data. BMC bioinformatics 18(1), 286.

Zmienko, A., Samelak, A., Kozlowski, P., and Figlerowicz, M. (2014). Copy number polymorphism in plant genomes. Theor Appl Genet 127(1), 1–18. doi: 10.1007/s00122-013-2177-7.

