## Supplementary Figure 1 for "SECNVs: A Simulator of Copy Number Variants and Whole-Exome Sequences from Reference Genomes"

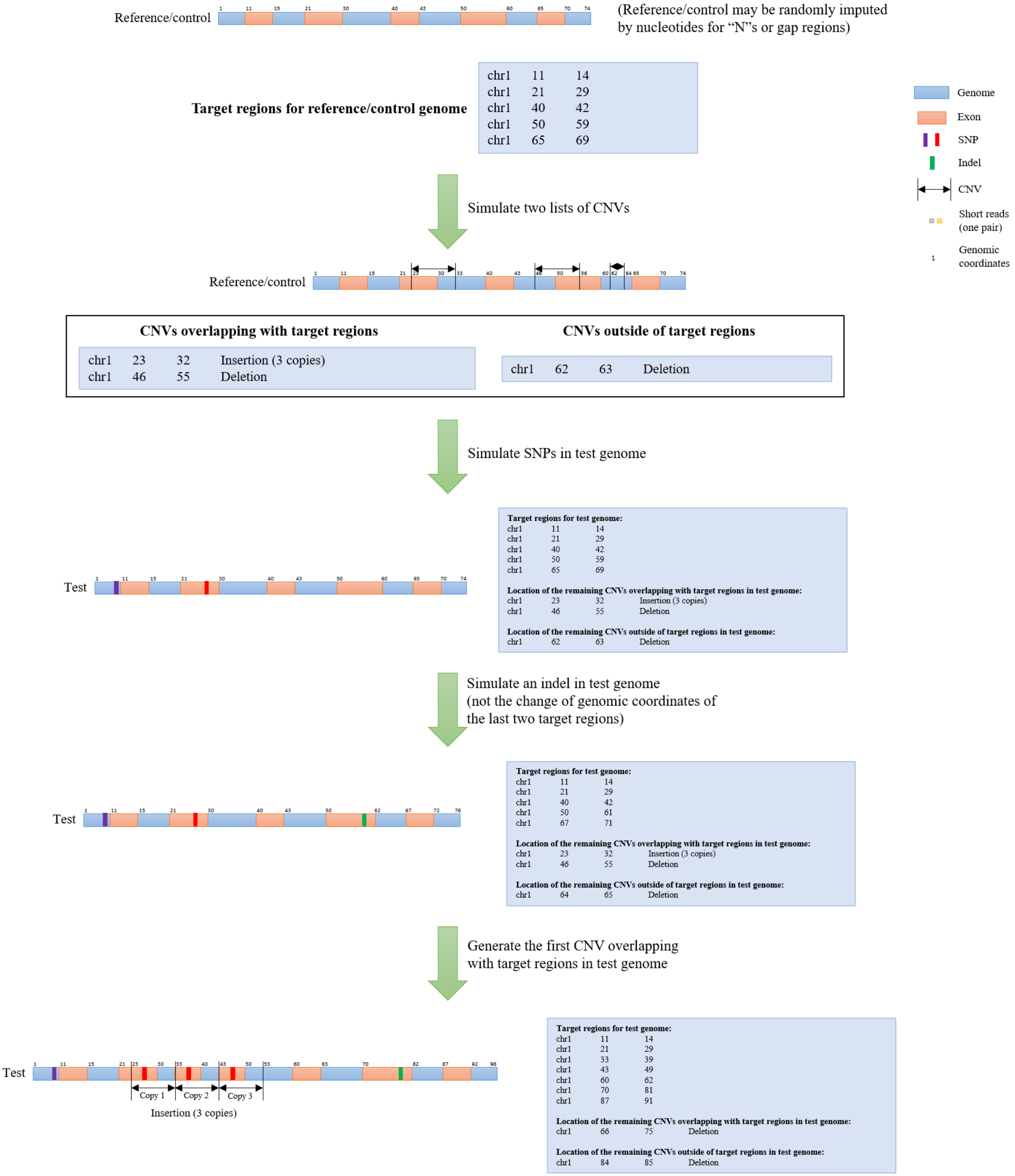
Reference/control
(Reference/control may be randomly imputed by nucleotides for “N”s or gap regions)
Target regions for reference/control genome
chr1 11 14
chr1 21 29
chr1 40 42
chr1 50 59
chr1 65 69
Genome
Exon
SNP
Indel
CNV
Short reads (one pair)
Genomic coordinates
Simulate two lists of CNVs
Reference/control
CNVs overlapping with target regions
chr1 23 32 Insertion (3 copies)
chr1 46 55 Deletion
CNVs outside of target regions
chr1 62 63 Deletion
Simulate SNPs in test genome
Test
Target regions for test genome:
chr1 11 14
chr1 21 29
chr1 40 42
chr1 50 59
chr1 65 69
Location of the remaining CNVs overlapping with target regions in test genome:
chr1 23 32 Insertion (3 copies)
chr1 46 55 Deletion
Location of the remaining CNVs outside of target regions in test genome:
chr1 62 63 Deletion
Simulate an indel in test genome
(not the change of genomic coordinates of the last two target regions)
Test
Target regions for test genome:
chr1 11 14
chr1 21 29
chr1 40 42
chr1 50 61
chr1 67 71
Location of the remaining CNVs overlapping with target regions in test genome:
chr1 23 32 Insertion (3 copies)
chr1 46 55 Deletion
Location of the remaining CNVs outside of target regions in test genome:
chr1 64 65 Deletion
Generate the first CNV overlapping with target regions in test genome
Test
Copy 1
Copy 2
Copy 3
Insertion (3 copies)
Target regions for test genome:
chr1 11 14
chr1 21 29
chr1 33 39
chr1 43 49
chr1 60 62
chr1 70 81
chr1 87 91
Location of the remaining CNVs overlapping with target regions in test genome:
chr1 66 75 Deletion
Location of the remaining CNVs outside of target regions in test genome:
chr1 84 85 Deletion

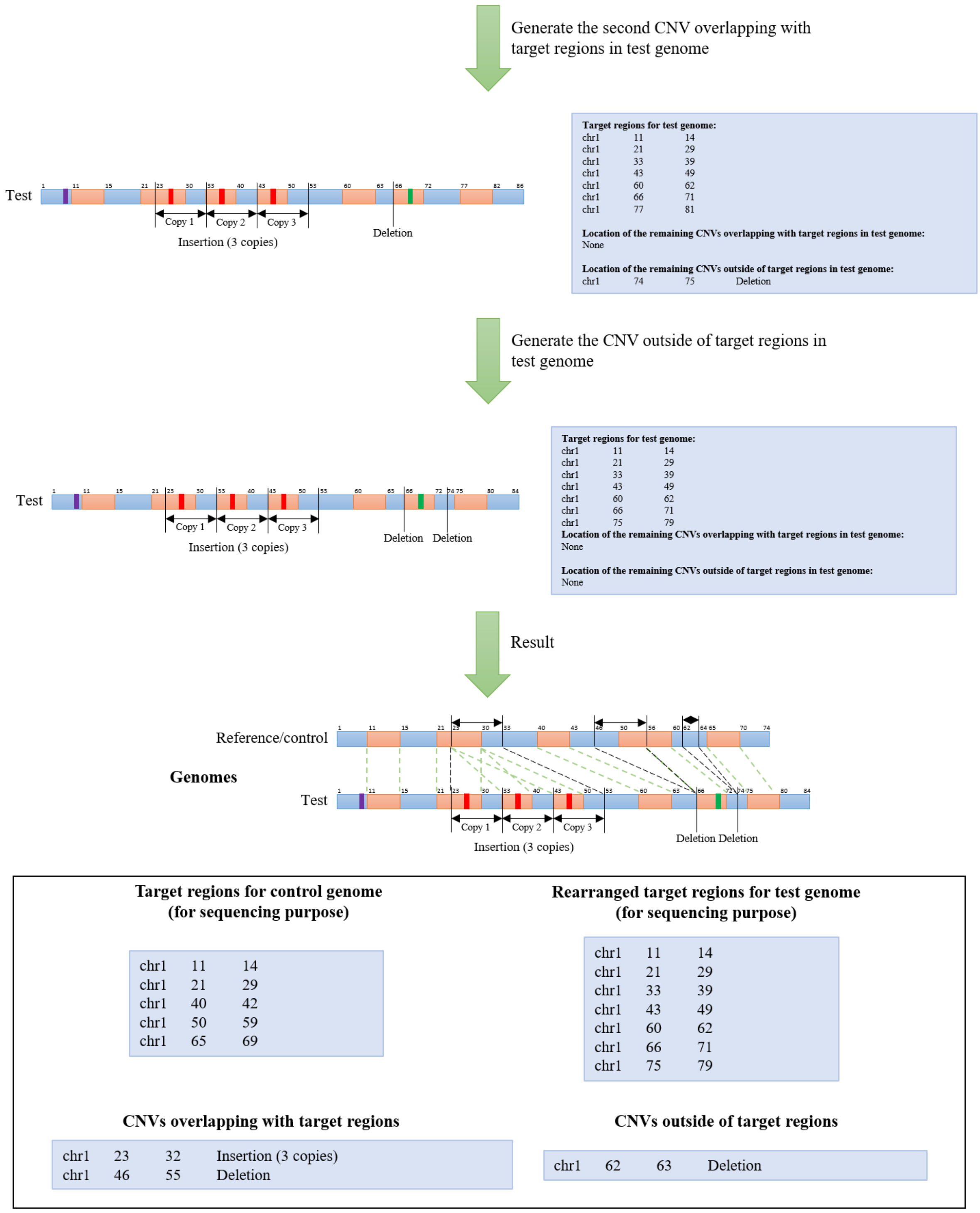


Short reads are then simulated for test and control genomes, and aligned back to the reference genome as shown in Figure 1. CNVs can then be detected by CNV detection tools.

**Supplementary Figure 1.** A small pseudo-genome was used as input to illustrate the simulation process and confirm that the code for the algorithm implemented in SECNVs is correctly simulating the test genome and target regions.
