## Supplementary File 1 for "SECNVs: A Simulator of Copy Number Variants and Whole-Exome Sequences from Reference Genomes"

**Supplementary File 1. Problem of Wessim.**

The approach used in Wessim1 is to cut the target regions into fragments based on the mean fragment size provided. However, Wessim1 only simulates reads at the start and end of target regions but it fails to generate fragments for simulated reads across the whole length of the target region. An example is shown at the target region chr1:134199214-134203590 in the mouse genome (mm10 assembly). Reads were only simulated at start and end of this target region but failed to be simulated across the whole target region. IGV was used for visualization.

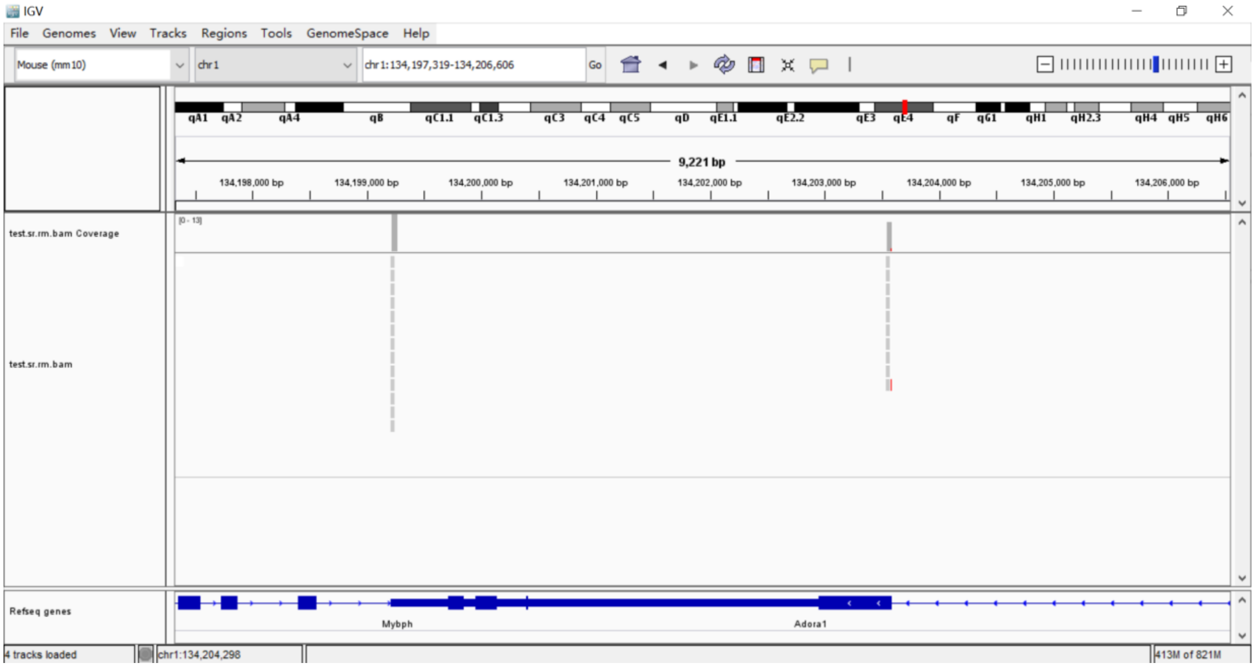
