## Supplementary File 2 for "SECNVs: A Simulator of Copy Number Variants and Whole-Exome Sequences from Reference Genomes"

**Supplementary File 2. Confirmation that SECNVs is correctly simulating the test genome and target regions.**

To validate the accuracy of rearranged genome and target regions, a mini genome was created as following (minig.fa):

>chr1
GNANNATCTCGTGNNNTGCACAANNNATATTGCATATCATGCGTNNCTATCANNNNNCATGGGGTGA
CCCGATC

Its target regions are as following (minit.bed):

| | | |
|---|---|---|
| chr1 | 11 | 14 |
| chr1 | 21 | 29 |
| chr1 | 40 | 42 |
| chr1 | 50 | 59 |
| chr1 | 65 | 69 |

Two CNV lists were supplied as input:

List of CNVs overlapping with target regions (mini_cnv_in.bed):

| chr | start | end | length | copy_number |
|---|---|---|---|---|
| chr1 | 23 | 32 | 10 | 3 |
| chr1 | 46 | 55 | 10 | 0 |

List of CNVs outside of target regions (mini_cnv_out.bed):

| chr | start | end | length | copy_number |
|---|---|---|---|---|
| chr1 | 62 | 63 | 2 | 0 |

For illustration purpose only, we manually made the 2 SNPs and indel like what is shown in Figure 3, because they would be otherwise generated randomly by SECNVs. Note that in SECNVs SNPs and indels were actually generated after random replacement of gap sequences.

Now the reference genome (minig_e.fa) was like:

>chr1
GNANNATCACGTGNNNTGCACAANNNTTATTGCATATCATGCGTNNCTATCANNNNNCCGATGGGGT
GACCCGATC

The 2 SNPs are at position 9 and 27 (purple and red). The indel is inserted at 59-60 (green).

And the target regions (minit_e.bed) were altered manually like:

| | | |
|---|---|---|
| chr1 | 11 | 14 |
| chr1 | 21 | 29 |
| chr1 | 40 | 42 |
| chr1 | 50 | 61 |
| chr1 | 67 | 71 |

List of CNVs overlapping with target regions (mini_cnv_in_e.bed) were not altered manually because they were before the indel:

| chr | start | end | length | copy_number |
|---|---|---|---|---|
| chr1 | 23 | 32 | 10 | 3 |
| chr1 | 46 | 55 | 10 | 0 |

List of CNVs outside of target regions (mini_cnv_out_e.bed) were altered manually because it was after the indel:

| chr | start | end | length | copy_number |
|---|---|---|---|---|
| chr1 | 64 | 65 | 2 | 0 |

These manually altered files were used as input. The command used was:

```
python SECNVs/SECNVs.py -G minig_e.fa -T minit_e.bed -rN gap -n_gap 3 \
-e_cnv mini_cnv_in_e.bed -o_cnv mini_cnv_out_e.bed
```

During the process, SECNVs randomly replaced gap sequences. A gap was defined as 3 or more consecutive "N"s. Now the reference genome was like:

>chr1
GNANNATCACGTGATCTGCACAAGACTTATTGCATATCATGCGTNNCTATCAGCTTACCGATGGGGTG
ACCCGATC

Which was proved by "control.fa":

```
>chr1
GNANNATCACGTGATCTGCACAAGACTTATTGCATATCATGCGTNNCTAT
CAGCTTACCGATGGGGTGACCCGATC
```

We didn't show the validation of excluding all "N" sequences or gap sequences here because CNV lists used as input were predefined, the parameter of excluding all "N" sequences would be ignored in this case.

The output target regions for test genome (the rearranged genome) were (test.target_regions_for_gen_short_reads.bed):

```
chr1    11    14
chr1    21    29
chr1    33    39
chr1    43    49
chr1    60    62
chr1    66    71
chr1    75    79
```

Which are in compliance with the anticipated results in Figure 3.

The output target regions for control genome were different than that of Figure 3 because we manually added an indel in the input genome (control.target_regions_for_gen_short_reads.bed), They were the same as in the manually altered "minit_e.bed":

```
chr1    11    14
chr1    21    29
chr1    40    42
chr1    50    61
chr1    67    71
```

The test genome (rearranged genome) should be:

>chr1
GNANNATCACGTGATCTGCACAAGACTTATTGAGACTTATTGAGACTTATTGCATATCATGCGTNTACC
GATGGTGACCCGATC

The 3-copy insertion was labeled blue. The bases before and after the deletions were labeled grey.

And was proved by “test.fa”:

```
>chr1
GNANNATCACGTGATCTGCACAAGACTTATTGAGACTTATTGAGACTTAT
TGCATATCATGCGTNTACCGATGGTGACCCGATC
```

To do everything similar to the example, but generating SNPs and indels randomly in the genome, the full command used would be:

```
python SECNVs/SECNVs.py -G minig.fa -T minit.bed -rN gap -n_gap 3 \
-s_r 0.04 -s_s 2 -i_r 0.025 -i_mlen 2 \
-e_cnv mini_cnv_in.bed -o_cnv mini_cnv_out.bed
```

We can also modify the above command to generate 0 CNVs by asking SECNVs to simulate a CNV larger than the genome size, and check its performance of making SNPs and indels:

```
python SECNVs/SECNVs.py -G minig.fa -T minit.bed -rN gap -n_gap 3 \
-s_r 0.04 -s_s 2 -i_r 0.025 -i_mlen 2 \
-e_tol 1 -min_len 1000
```

The output control genome was:

>chr1
GNANNATCACGTGAAGTGCACAAGTCTTATTGCATATCATGCGTNNCTATCATTCAGCCGATGGGGTG
ACCCGATC

```
>chr1
GNANNATCACGTGAAGTGCACAAGTCTTATTGCATATCATGCGTNNCTAT
CATTCAGCCGATGGGGTGACCCGATC
```

The output test genome was:

>chr1
GNANNATCACGTGAAGTGCGCAAGTCTTATTGCATATCATGCGTNNCTATCATTCGGCCGATGGGGCT
GACCCGATC

```
>chr1
GNANNATCACGTGAAGTGCGCAAGTCTTATTGCATATCATGCGTNNCTAT
CATTCGGCCGATGGGGCTGACCCGATC
```

The 2 SNPs were labeled red and the indel with length <= 2 was labeled green.
